# Transcription factor retention through multiple polyploidisation steps in wheat

**DOI:** 10.1101/2022.02.15.480382

**Authors:** Catherine EB Evans, Ramesh Arunkumar, Philippa Borrill

## Abstract

- Whole genome duplication (WGD) is widespread in plant evolutionary history, but the mechanisms of non-random gene loss after WGD are debated. The gene balance hypothesis proposes that dosage-sensitive genes such as regulatory genes are retained in polyploids. To test this hypothesis, we analysed the retention of transcription factors (TFs) in the recent allohexaploid bread wheat *(Triticum aestivum*).
- We annotated TFs in hexaploid, tetraploid and diploid wheats; compared the co-expression of homoeologous TF and non-TF triads; and analysed single nucleotide variation in TFs across cultivars.
- We found that, following each of two hybridisation and WGD events, the proportion of TFs in the genome increased. TFs were preferentially retained over other genes as homoeologous groups in tetraploid and hexaploid wheat. Across cultivars, TF triads contain fewer deleterious missense mutations than non-TFs.
- TFs are preferentially retained as three functional homoeologs in hexaploid wheat, in support of the gene balance hypothesis. High co-expression between TF homoeologs suggests that neo- and sub-functionalisation are not major drivers of TF retention in this young polyploid. Knocking out one TF homoeolog to alter gene dosage, using TILLING or CRISPR, could be a way to further test the gene balance hypothesis and generate new phenotypes for wheat breeding.

## Introduction

Gene duplication plays a major role in the evolution of genetic and phenotypic diversity and in speciation events in eukaryotes (Lynch & Conery, 2000; Van de Peer *et al*., 2009b). Ancient whole genome duplications (WGD) are observed throughout the angiosperm plant phylogeny and occurred at the base of major clades such as the seed plants, core eudicots and monocots (Van de Peer *et al*., 2009a; Tasdighian *et al*., 2017). Following WGD, most gene duplicates are eventually lost from the genome by pseudogenisation or deletion over the course of millions of years (Freeling, 2009), although the retention of a significant portion of duplicates has been observed across multiple plant lineages (Lloyd *et al*., 2014; Li *et al*., 2016).

Several explanations for the retention of gene duplicates have been proposed including selection for genetic redundancy (or genetic buffering) where effects of a null mutation are compensated by the presence of an intact duplicate copy (Nowak *et al*., 1997) and gene dosage increase where increased dosage is advantageous, for example by increasing flux through a pathway (Ohno, 1970). Other mechanisms include sub-functionalisation of the duplicate copies which may be mediated by complementary degenerative mutations in each copy (Force *et al*., 1999) and paralog interference where degenerative mutations in one duplicate copy interfere with the function of the other copy thereby promoting the retention of both functional copies (Baker *et al*., 2013). Duplicate gene copies may also be retained if they evolve new and distinct functions through neo-functionalisation (Ohno, 1970).

An alternative model for duplicate retention is the gene balance hypothesis, which proposes that dosage sensitive genes tend to be retained as duplicates (Birchler *et al*., 2005; Birchler & Veitia, 2007). This hypothesis explains the observation that the loss of genes after WGD is non-random and certain classes of gene are preferentially retained including genes involved in regulatory interactions or in protein complexes which are dosage sensitive (Blanc & Wolfe, 2004). Conversely, these dosage sensitive genes are less frequently found in segmental duplications in which they would upset the dosage balance with interacting partners (Maere *et al*., 2005), in contrast to WGD where their interacting partners would also be duplicated.

Studies across multiple angiosperms have revealed that transcription factors (TFs), a major type of dosage sensitive regulatory gene, tend to be retained as duplicates after WGD for millions of years (Lloyd *et al*., 2014; Li *et al*., 2016). In a comparative study of 37 sequenced angiosperm genomes, Li *et al*. (2016) found that duplicate genes that originated at the Cretaceous-Paleogene boundary ~50 to 70 million years ago (mya) when a large number of WGD events occurred (Van de Peer *et al*., 2009a), were enriched for TFs. However these angiosperm-wide studies focussed on relatively old WGD events >5mya, whilst more recent WGD events which have occurred in individual lineages and are found in several major crop species are less well studied, perhaps due to a lack of genome sequences. Preferential retention of dosage sensitive genes such as TFs has been observed in the young polyploid *Tragopogon miscellus* which underwent WGD only ~80 years before (Buggs *et al*., 2012). This study used a limited number of loci, therefore there remains a need to understand the effects of recent (<5 mya) WGD at a genome-wide scale. The recent publication of the wheat genome sequence (IWGSC *et al*., 2018) provides an opportunity to examine the retention of dosage-sensitive genes from more recent WGDs.

Hexaploid bread wheat evolved from two hybridisation and WGD events: allotetraploid wild emmer wheat (*Triticum turgidum* ssp. *dicoccoides*) was formed approximately 0.4 mya when the A genome progenitor *Triticum urartu* hybridised with the B genome progenitor species (Feldman & Levy, 2012). The allotetraploid emmer was domesticated and hybridised with the D genome progenitor *Aegilops tauschii* approximately 10,000 years ago to form hexaploid bread wheat (*Triticum aestivum* L.) (Dubcovsky & Dvorak, 2007). This two-step recent history of WGD events has resulted in >50% of genes being present with three homoeologous copies in bread wheat (IWGSC *et al*., 2018). Previous studies in wheat have shown that 58% of NAC TFs and 63% of MIKC-type MADS-box TFs have three homoeologs (Borrill *et al*., 2017; Schilling *et al*., 2020), but a systematic study has not been carried out to establish whether the preferential retention of TFs is observed across all TF families in this recent polyploidy.

In this study we investigated whether these two recent WGD events resulted in the preferential retention of TFs in hexaploid wheat, as would be predicted by the gene balance hypothesis. Using the extensive curated expression data available for wheat, we explored alternative hypotheses about TF retention, such as sub- or neo-functionalisation, based on expression patterns. Moreover, since genetic variation in several TFs has been instrumental in wheat adaptation during domestication including the free-threshing gene *Q* (Simons *et al*., 2006) and the vernalisation gene *VRN1* (Yan *et al*., 2003), we examined the natural variation in TF homoeologs observed in wheat. Specifically we examined the propensity of wheat TFs to be retained as functional copies without deleterious mutations at a population level. Hence our study addresses not only an evolutionary question about the retention of TFs in young polyploids, but also provides insight into TF expression diversity and genetic variation which lays a foundation for future research and breeding.

## Materials and Methods

### Annotation of TFs in wheat and progenitor species

Peptide sequences for genes in the RefSeqv1.1 gene annotation of *Triticum aestivum* cv. Chinese Spring (IWGSC *et al*., 2018) were downloaded from EnsemblPlants (Howe *et al*., 2020). The file was divided into three parts to contain <50,000 sequences per file and TFs were annotated in each file using iTAK online v1.6 (Zheng *et al*., 2016). In cases where different transcript isoforms were assigned to different TF families (23 out of 6,128 genes) the family assigned to the longer transcript isoform was retained (Table S1). Peptide sequences for genes in the *Aegilops tauschii* assembly (Luo *et al*., 2017) were downloaded from EnsemblPlants, divided into six smaller files and annotated using iTAK online v1.6. Again, when different transcript isoforms were assigned to different TF families (186 out of 2,120 genes) the family assigned to the longer transcript isoform was retained (Table S2). In general, discrepancies between TF families were due to one isoform being truncated, with the truncated isoform lacking a protein domain that allowed a more specific TF family to be assigned to the longer isoform. Coding sequences for the longest isoforms of genes in the *Triticum urartu* genome (Ling *et al*., 2018) were downloaded from http://www.mbkbase.org/Tu/ and annotated using iTAK online v1.6 (Table S3). TF annotations for *Triticum turgidum* ssp. *dicoccoides* cv. Zavitan (Avni *et al*., 2017) were downloaded from the iTAK database (update 18.12) (Zheng *et al*., 2016) (Table S4).

### Identification of 1:1:1 triads in hexaploid and 1:1 diads in tetraploid wheat

Homoeologs were downloaded from EnsemblPlants Biomart for the RefSeqv1.1 gene annotation using only high confidence gene models. Only one2one homoeologs (assigned by EnsemblPlants) were retained. There were 20,393 triads corresponding to 61,179 genes (56.7% of genes) (Table S1). Homoeologs in *T. turgidum* ssp. *dicoccoides* were obtained from Avni *et al*. (2017) and filtered to only retain 1:1 homoeologs by removing “singleton” and “hit2homolog” (i.e. paralog) groups (Table S4). Only high confidence genes from the RefSeqv1.1 annotation were used in all subsequent analyses.

### Adjusting for the effect of gene loss in tetraploid wheat on hexaploid wheat triad numbers per TF family

In order to adjust for the differences in triad proportions between TF families observed in hexaploid due to the varying proportions in diads in tetraploid wheat, we calculated the normalised percentage of genes in triads:

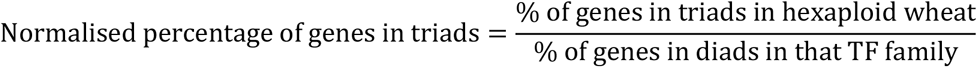

For example if 60% of genes were in triads in hexaploid, but only 80% genes were in diads in tetraploid, the normalised value will be 75% - i.e. 75% of the potential triads were formed because we have accounted for the 20% which were already missing in tetraploid.

### Correlation of expression levels per family to homoeolog retention in triads

To measure the gene expression level of each TF family we used RNA-seq data from 15 different tissues and developmental stages from Chinese Spring (Choulet *et al*., 2014). These included tissues from seedling roots and shoots through to grain 30 days after anthesis. We downloaded gene expression data in transcripts per million (tpm) for this dataset from expVIP (www.wheat-expression.com) (Borrill *et al*., 2016; Ramírez-González *et al*., 2018). We calculated the mean expression level for each gene across the 15 tissues, and then calculated the median expression level for each TF family. We fitted a linear regression model between log(median expression level per TF family) and the percentage of TFs in triads in the family.

### Correlation of tandem duplication per family to homoeolog retention in triads

For all TF genes we defined tandem duplicates as genes which were adjacent in the genome assembly according to their gene IDs ± 3 genes in either direction (gene IDs increase by 100 for adjacent genes in this genome assembly). We allowed 1 or 2 genes between tandem duplicates because a tandem duplication event may have occurred capturing a TF and non-TF in the same duplication event. Each nearby duplicate was counted as one tandem duplication event (i.e. a cluster of 3 TF genes would be counted as 2 tandem duplication events), and the total number of tandem duplication events was divided by the total number of genes in each TF family to calculate the percentage of tandem duplicated genes per TF family. We fitted a linear regression model between the percentage of genes which are tandem duplicates per TF family and the percentage of TFs in triads in the family. We repeated our analysis only considering ±2 genes (with one gene between them) or ±1 gene (with no gene between them) as tandem duplicates.

### Calculation of homoeolog similarity of expression per family

Using the same data from 15 different tissues and developmental stages from Chinese Spring we filtered to only keep triads where at least 1 homoeolog was expressed >0.5 tpm in one tissue (calculated as the mean value of two biological replicates). To account for differences in expression level between TFs and non-TFs, we normalised the expression level of each triad per tissue to sum to 1 as in Ramírez-González *et al*. (2018) before calculating the standard deviation of expression level between homoeologs. For 58 out of 19,391 triads (0.3%) the TF family was inconsistent between homoeologs (e.g. MYB and MYB-related) so the family assigned to two of the three homoeologs was retained. A Mann-Whitney test was used to determine whether the standard deviation within TF triads was different from non-TF triads for each tissue.

### Calculation of homoeolog co-expression per family

To calculate the Pearson’s correlation between the three homoeologs we used the same data from 15 different tissues and developmental stages from Chinese Spring. We filtered to only keep triads where at least 1 homoeolog was expressed >0.5 tpm in one tissue (calculated as the mean value of two biological replicates), and triads where all three homoeologs were expressed (tpm>0 in at least one tissue). The Pearson’s correlation was calculated between homoeologs within a triad in a pairwise fashion (A vs B, B vs D, A vs D) and the three correlations were plotted for each triad. To calculate the median Pearson’s correlation for TF triads and non-TF triads, the Pearson’s correlation values were Z transformed using DescTools v0.99.44 (Signorell, 2021) before obtaining the median, then back-transformed to reduce bias (Corey *et al*., 1998).

As an alternative measure of co-expression we used information about module assignment from a Weight Gene Co-expression Network Analysis (WGCNA) across 850 wheat RNA-samples (Langfelder & Horvath, 2008; Ramírez-González *et al*., 2018). The co-expression network was built using RefSeq v1.0 annotation. To enable compatibility with our TF annotation which was carried out using RefSeq v1.1 annotation, only genes which were 99% identical with >90% coverage from v1.0 to v1.1 were included in this analysis. To calculate the percentage of triads with homoeologs in the same module only triads in which all three homoeologs had a module assigned, excluding module 0, were considered. Module 0 largely contains genes with invariable expression patterns between samples (Ramírez-González *et al*., 2018).

### Analysis of SNP variation data

To investigate the types of single nucleotide polymorphisms (SNPs) in wheat TFs, we used exome capture data of 811 hexaploid wheat landraces and cultivars representing global genetic diversity (He *et al*., 2019). Filtered and imputed SNPs (~3 million) were downloaded May 2021 from http://wheatgenomics.plantpath.ksu.edu/1000EC/.

We selected SNPs in genes in triads and used the Ensembl Variant Effect Predictor (VEP v99.2) to predict the effect of SNPs on these genes (McLaren *et al*., 2016). From an input of 529,066 SNPs in triad genes, VEP output 1,146,195 SNP effects. We selected 216,285 SNPs predicted in the coding sequence of the canonical transcript of a triad gene. Using R, we filtered to exclude: SNPs which were also splice region variants; missense variants without SIFT scores; and SNPs with >25% missing calls. 210,578 SNPs remained (97% of unfiltered SNPs in coding sequences of canonical transcripts).

To exclude potential bias from rare SNPs, we filtered to retain SNPs with a minor allele frequency (MAF) of at least 0.01, resulting in a total of 74,442 SNPs. To focus on SNPs more likely to have a functional effect *in planta*, we only retained SNPs in genes that were expressed at >0.5 tpm in at least one tissue using data from (Choulet *et al*., 2014). We excluded SNPs in regions that He *et al*. (2019) identified as being under environmental adaptation, improvement selection or within a selective sweep, as positive and purifying selection have similar impacts on nucleotide diversities in populations (Cvijovic *et al*., 2018). Introgressed sites were also excluded as they would have had a different demographic history compared to the remainder of the genome. Synonymous sites that had more than one annotation were excluded from analyses. This left 16,119 SNPs (1020 TF, 15,099 non-TF).

We categorised the SNPs according to variant effect (stop gained, missense and synonymous). Missense mutations were further categorised as deleterious or tolerated according to their Sorting Intolerant from Tolerant (SIFT) prediction (Sim *et al*., 2012). A SIFT score of ≤0.05 is predicted to be deleterious, affecting the protein phenotype and a score >0.05 is predicted to be tolerated, not affecting phenotype.

Per site nucleotide diversity was estimated using VCFtools c0.1.16 (Danecek *et al*., 2011). Mann-Whitney tests were used to compare the TF and non-TF nucleotide site diversity distributions. Mutation load was estimated by calculating the number of homozygous alternate alleles for each site type, divided by the summed lengths of all the canonical transcripts for TFs and non-TFs separately. A linear regression with mutation load as the response and the category of sites (stop gained, deleterious missense, tolerated missense, synonymous) and the group of genes (TF and non-TF) was fitted, and an ANOVA was performed to test for the significance of the fixed effects. Further, a Tukey’s test was used to compare TFs and non-TFs for each site category. Individuals with extreme mutation loads were classed as those with loads in the 2.5% tails in any of the distributions. For the TF families plot, we excluded SNPs which are only represented in individuals with extreme mutation loads. We plotted the proportion of SNPs by variant effect for TF families containing more than 10 triads and ≥5 SNPs and for non-TFs.

## Results

### TFs homoeologs are retained across polyploidisation events more frequently than non-TFs

To explore TF evolution and conservation in polyploid wheat we annotated TFs in the hexaploid *T. aestivum* (AABBDD), the tetraploid ancestor *T. turgidum* ssp. *dicoccoides* (AABB) and the diploid ancestral species *T. urartu* (AA) and *A. tauschii* (DD) (Table S1-4). We found that overall the percentage of genes in the genome which were annotated as TFs increased as the ploidy level increased (Figure 1a), from 4.4% in diploid *T. urartu*, to 4.9% in tetraploid *T. turgidum* ssp *dicoccoides*, to 5.7% in hexaploid *T. aestivum*. A higher percentage of genes were TFs in *A. tauschii* (5.4%) than in the other diploid progenitor *T. urartu* (4.4%), although this was still lower than in the hexaploid wheat (5.7%). This supports the hypothesis that TFs are preferentially retained, compared to other types of genes, in polyploid wheat. The retained TFs were distributed similarly across the genomes in tetraploid (50.4% on A genome, 49.6% on B genome) and hexaploid wheat (33.7% on A genome, 33.1% on B genome and 33.3% on D genome), consistent with previous reports that wheat does not show biased sub-genome fractionation associated with preferential loss of genes associated with one subgenome (IWGSC, 2014).

**Figure 1.**
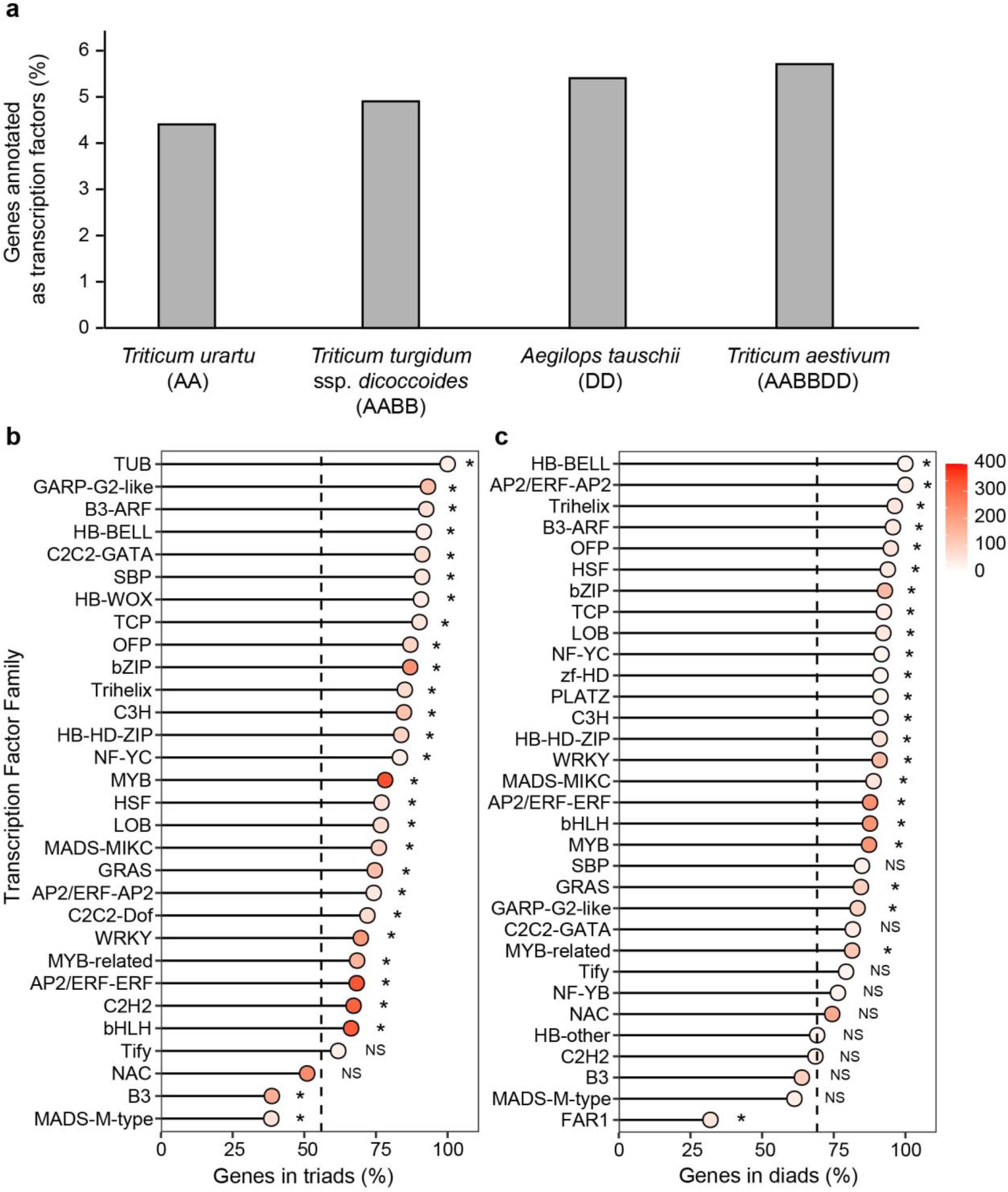
Transcription factor (TF) genes in *T. aestivum* and ancestral species. a) Percentage of genes annotated as TFs in hexaploid *T. aestivum* and the tetraploid and diploid ancestral species. b) Percentage of genes in triads in *T. aestivum* TF families with >10 triads and c) Percentage of genes in diads in *T. turgidum* ssp. *dicoccoides* TF families with >10 diads. In b) and c) the dotted black line indicates the mean value for non-transcription factors and asterisks (*) denote families which are significantly different from non-TFs (Fisher’s exact test, p<0.05, FDR corrected for multiple testing). NS= non-significant. The fill colour of the dots indicates the number of genes in the TF family.

We hypothesised that the higher proportion of TF genes in polyploid wheats compared to their wheat progenitors were due to the preferential retention of TF homoeologs, whilst other types of genes were less often retained with all homoeologs. Consistent with this hypothesis we found that in polyploid wheat, TFs were more frequently present with all homoeologs than other types of genes. Across TF and non-TF genes in hexaploid *T. aestivum*, 56.7% of genes are in triads with a single A homoeolog, a single B homoeolog and a single D homoeolog. TF genes were more commonly found in triads with 70.5% of TFs in triads, compared to other types of genes (55.9% in triads; p<0.001, Fisher’s exact test). This enrichment for triads was observed in nearly all TF families (Fig 1b, Fig S1). Similar trends were observed in tetraploid *T. turgidum* ssp. *dicoccoides*. Across TF and non-TF genes 69.8% of genes in the tetraploid were in diads with a single A homoeolog and a single B homoeolog, but this figure rose to 82.5% of TFs, compared to 69.2% of other types of genes (p<0.001, Fisher’s exact test). The enrichment for diads was common to most TF families (Figure 1c, Fig S2).

In general, TF families with a lower percentage of triads in hexaploid wheat already had a lower proportion of diads in tetraploid. For example, the B3 and MADS-M-type families had fewer triads/diads in both wheat species than non-TF genes, with tetraploid having 63.9% and 61.3% of genes for the B3 and MADS-M-type family in diads respectively, and hexaploid having 38.7% and 38.5% of genes in triads respectively (Figure 1b and 1c). The NAC TF family, which is one of the largest TF families in wheat, is one of the less well retained TF families in tetraploid (74.5% of genes in diads), although this is still higher than for non-TFs. However, in hexaploid wheat only 51.0% of NACs are in triads which is lower than for non-TFs. After accounting for gene loss in tetraploid wheat, the B3, MADS-M-type and NAC families in hexaploid wheat still had significantly fewer genes in triads (60.5%, 62.8% and 68.5% respectively) than non-TFs (80.8%; FDR adjusted p<0.001 Fisher’s exact test). This indicates that homoeolog loss in specific TF families occurred across both polyploidisation steps and was not solely due to pre-existing gene loss in the tetraploid.

### Differential conservation of TF families as triads is correlated with expression level and tandem duplications

To understand why certain TF families are more prone to homoeolog loss we explored two previously proposed hypotheses. The first is the “highly expressed gene retention idea” (Freeling, 2009), which proposes that gene families which are highly expressed are more likely to be retained with homoeologous copies. Secondly we investigated the “balanced gene drive hypothesis” (Freeling, 2009) which proposes that gene families which have more tandem duplications are less likely to be retained with homoeologous copies.

To test the correlation between gene expression level and gene retention in hexaploid wheat we used RNA-seq data from 15 different tissues from Chinese Spring from a developmental timecourse (Choulet *et al*., 2014). We calculated the mean expression level for each gene across the 15 tissues, and then calculated the median expression level for each TF family. Focussing on TF families with >10 triads we found a significant positive correlation between the expression level of the TF family and the percentage of genes in the TF family which are in triads (R^2^ = 0.40, p<0.001; Figure 2a). This relationship also held across all TF families regardless of size, although the correlation was weaker due to small families that were outliers (R^2^ = 0.21, p<0.001; Fig S3). Consistent with this relationship, the three TF families with a lower retention of homoeologs in hexaploid wheat than non-TFs (NAC, MADS-M-type and B3) all had low median expression levels (Figure 2a).

**Figure 2.**
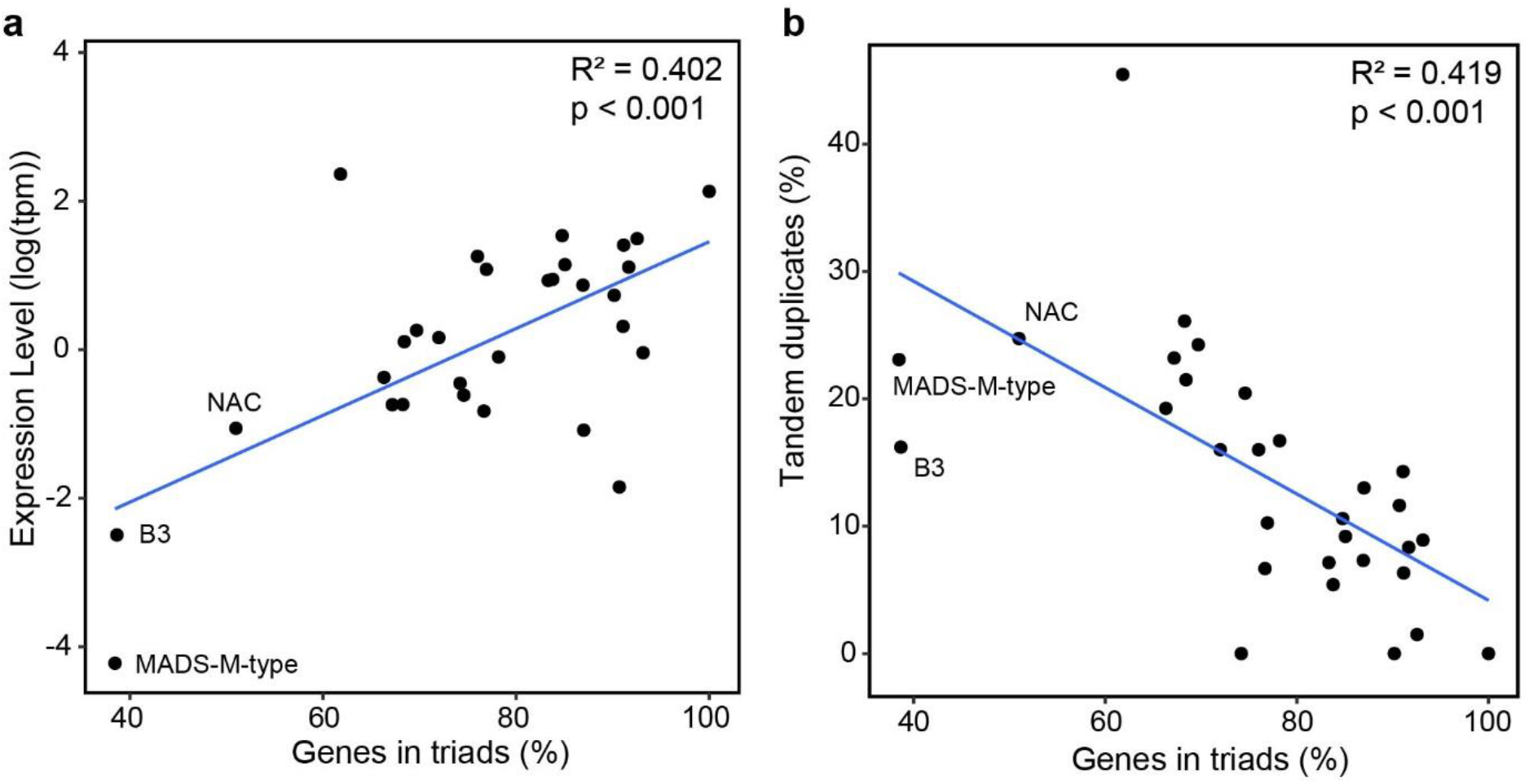
Factors explaining differential retention of homoeologs in different transcription factor (TF) families. a) Median expression level per TF family plotted against the percentage of the TF family in triads for TF families with >10 triads. The mean expression level of each gene in transcripts per million (tpm) was calculated using 15 tissues of Chinese Spring RNA-seq data and these gene level values were used to calculate median expression level within the TF family. b) The percentage of tandem duplicated genes within each TF family plotted against the percentage of the TF family in triads for TF families with >10 triads. TFs were considered to be tandem duplicates when they were up to ±3 genes away from each other (i.e. up to two genes in between duplicates).

We also explored the relationship between tandem duplications and gene retention. Focussing on TF families with >10 triads we found that the degree of tandem duplication in a TF family was negatively correlated with the percent of triads within the TF family, consistent with the “balanced gene drive hypothesis” (R^2^ = 0.42, p<0.001, permitting up to 2 genes between tandem duplicated TFs; Figure 2B). This correlation held with a more stringent criteria for tandem duplicates only permitting 1 gene between tandem duplicates (R^2^ = 0.38, p<0.001; Fig S4a) or 0 genes between tandem duplicates (R^2^ = 0.28, p=0.003; Fig S4B). These relationships also held when including all TF families regardless of size, although the correlation was weaker (R^2^ = 0.14 to 0.22, p < 0.003) due to variability within small families (Fig S4c-e). The NAC TF family that had low retention of homoeologs in hexaploid wheat had quite high levels of tandem duplication (Figure 2b). However, the MADS-M-type and B3 TF families had lower levels of tandem duplication than the trendline across all TF families (Figure 2b), suggesting that low expression levels (Figure 2a) may be driving the lack of homoeolog retention in these families. Together these results indicate that different retention levels in individual TF families are associated with gene expression level and the degree of tandem duplication.

### TF triads do not show increased sub- or neo-functionalisation of expression or co-expression patterns

There are several different mechanisms which can contribute to the retention of homoeologs following polyploidisation. Conant *et al*. (2014) proposed a pluralist framework in which dosage effects, sub-functionalisation and neo-functionalisation interplay to preserve duplicated genes, in a time dependent manner. Although transcriptomics cannot provide a definitive answer about the contributions of these different mechanisms (Conant *et al*., 2014), it can provide a starting point to understand potential mechanisms operating.

First, we used the same RNA-seq samples from 15 tissues from Chinese Spring to test whether TF triads had more different expression levels between homoeologs than non-TF triads, which would support sub- or neo-functionalisation of TF homoeologs at the gene expression level leading to TF retention. We normalised the global expression of each triad so that total expression level of the triad was 1 as described in (Ramírez-González *et al*., 2018), to account for differences in expression level between TFs and non TFs. We found that the standard deviation between the expression levels of homoeologs within TF triads was not significantly different from non-TF triads in 14 out of 15 tissues (Mann-Whitney test, p>0.05). Only roots at Zadoks stage 39 (flag leaf ligule just visible) had a significantly lower standard deviation between homoeolog expression levels in TF triads than in non-TF triads (median 0.093 for TF triads, 0.099 for non-TF triads, p = 0.036, Mann-Whitney test). Overall, the standard deviation between homoeolog expression levels was not higher in TFs than non-TFs in any tissue suggesting that sub- or neo-functionalisation is not occurring at the gene expression level between TF homoeologs globally.

Building upon this finding, we explored co-expression between homoeologs across different tissues. We calculated the Pearson’s correlation coefficient pairwise between homoeologs across the 15 Chinese Spring tissues. Co-expression was higher for TF triads than non-TF triads (Pearson’s correlation coefficient 0.938 vs 0.923, p-value <0.001, Mann-Whitney test). Amongst TF families with over 10 triads, most TF families showed higher homoeolog co-expression than non-TFs, and the differences were significant for nine TF families (Figure 3a). Two TF families has significantly lower homoeolog co-expression than non-TFs (OFP and MADS-M-type, Figure 3a) The trend for higher co-expression within TF families than non-TFs was also observed in TF families with fewer than 10 triads (Fig S5).

**Figure 3.**
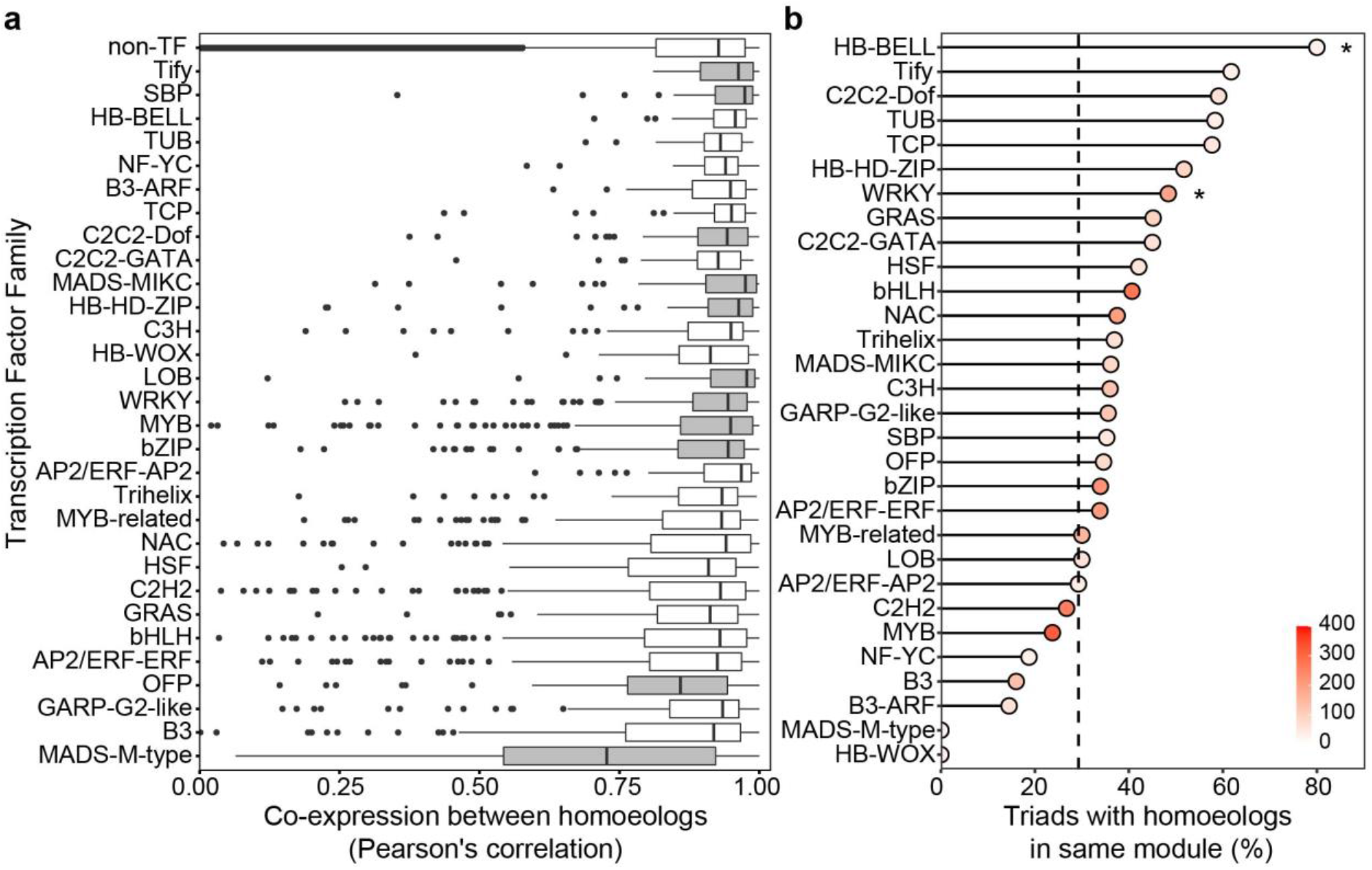
Co-expression of homoeologs within triads in transcription factor (TF) families with >10 triads. a) Pearson’s correlation coefficient between homoeologs across 15 tissues per TF family. TF families which were significantly different to non-TFs are highlighted in grey (Mann-Whitney test, p<0.05, FDR corrected for multiple testing). The correlation between non-TF homoeologs is shown in the top row. b) Homoeologs in same module in 850 sample WGCNA network per TF family. Black dotted line in b) represents mean value of non-TFs and asterisks (*) denote families which are statistically significant different from non-TFs (Fisher’s exact test, p<0.05, FDR corrected for multiple testing). The fill colour of the dots in b) indicates the number of genes in the TF family.

As an alternative measure of triad co-expression we explored a previously generated co-expression network made using WGCNA across 850 wheat RNA-samples (Langfelder & Horvath, 2008; Ramírez-González *et al*., 2018). We found that TF homoeologs were more frequently assigned to the same co-expression module than non-TF homoeologs (35.5% vs 29.3%; p<0.001 Fisher’s exact test), consistent with our Pearson’s correlation approach. A higher level of co-expression in TFs than non-TFs was consistent across most TF families in this WGCNA based approach although the difference was only statistically significant in a few families after adjustment for multiple testing (Figure 3b and Fig S6). TF families which showed higher co-expression were quite consistent with both measures of co-expression, e.g. Tify and WRKY, whilst some other families such as MADS-M-type TFs had lower co-expression using both measures (Figure 3). Overall, we did not find support for higher levels of sub- or neo-functionalisation at the expression or co-expression level in TF triads than in non-TFs, suggesting that other mechanisms such as dosage may be important for TF retention.

### Reduced deleterious mutation load in TF triads compared to non-TFs

To investigate how TFs evolve in wheat populations we explored single nucleotide polymorphisms (SNPs) in TFs and non-TFs using an exome capture dataset of 811 diverse hexaploid wheat cultivars and landraces (He *et al*., 2019). We hypothesised that TF triads would accumulate fewer mutations deleterious to gene function than non-TF triads, which would be consistent with their preferential retention during polyploidisation. We did not observe significant differences in the distribution of deleterious or synonymous nucleotide site diversities, estimated using π, between TFs and non-TFs (Fig S7). π is low when allele frequency is low or high (Fig S8) and, therefore, it does not capture the deleterious load burden in TFs and non-TFs. To identify the mutational burden, we calculated the number of homozygous deleterious and synonymous mutations in TF and non-TF triads. Numbers of homozygous mutations per individual scaled by the total length of all canonical transcripts differ between TF and non-TF genes (ANOVA, F=66.5, df=1, p<0.001). There were 32.0% fewer deleterious missense mutations per kilobase in TFs compared to non-TFs (Figure 4a; p<0.001, Tukey’s test). Frequencies of homozygous stop gained mutation were not significantly different between TFs and non-TFs. However, only 7 stop gained mutations were detected in TFs making the comparison underpowered. There were 5.7% more tolerated missense mutations and 17.6% fewer synonymous mutations per kilobase in TFs compared to non-TF genes. As sites occurring in regions associated with adaptation, introgression, or domestication were removed, the lower synonymous site diversity and load in TFs likely reflects background selection.

**Figure 4.**
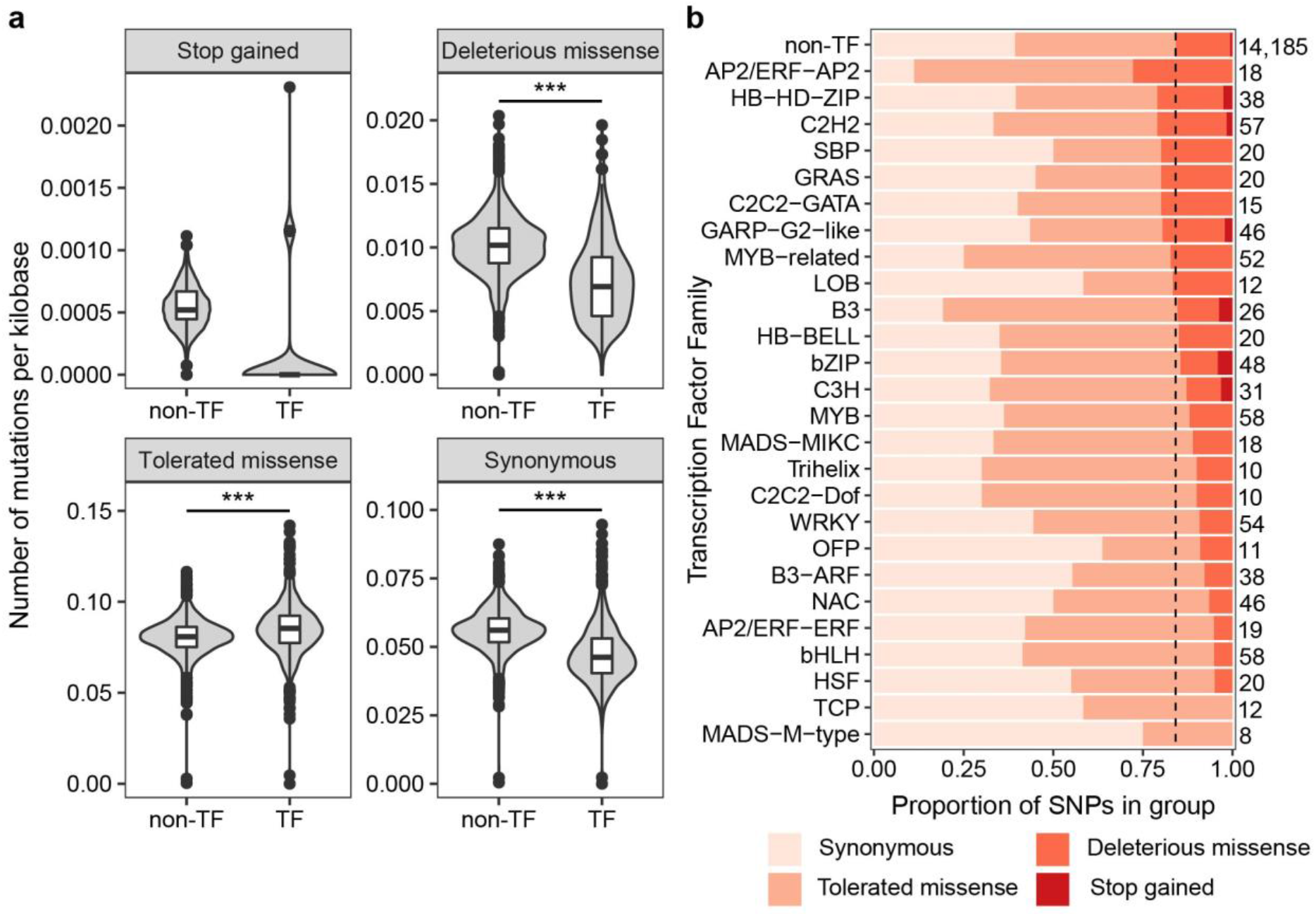
Transcription factors (TFs) accumulate fewer harmful mutations than non-TFs. a) Mutational burden in TFs compared to non-TFs for different categories of variants in 811 individuals. Mutational burden was calculated as the number of homozygous mutations of each type per individual scaled by the total length of all canonical transcripts for TFs and non-TFs. The total number of polymorphic sites analysed for TFs and non-TFs were: stop gained (TF = 7, non-TF = 107), deleterious missense (TF = 125, non-TF = 2,277), tolerated missense (TF = 474, non-TF = 6,723), synonymous (TF = 414, non-TF = 5,989). *** indicates p < 0.001 following a Tukey’s test comparing the different classes of variants. b) The proportion of single nucleotide polymorphisms (SNPs) by variant effect in TF families containing > 10 triads and ≥ 5 SNPs. These SNPs are in genes expressed in at least one tissue and have minor allele frequency ≥ 0.01. The number of SNPs in each group is shown to the right of the bars. TF families are sorted according to the proportion of deleterious missense plus stop gained SNPs, and non-TFs are shown in the top row. The black dotted line represents the split between (synonymous + tolerated missense) and (deleterious missense + stop gained) SNPs in non-TFs.

To explore the distribution of SNP effects across TF families, we plotted the proportion of SNPs of different effects in the coding sequence of TFs in families containing >10 triads and ≥5 SNPs (Figure 4b). 17 out of 26 TF families had fewer deleterious missense plus stop gained SNPs relative to non-TFs, while 9 had more. The lowest proportion of deleterious plus stop gained SNPs were found in the MADS-M-Type (0.0%), TCP (0.0%) and HSF (5.0%) families, and the highest proportion in the AP2/ERF-AP2 (27.8%), HB-HD-ZIP (21.1%) and C2H2 (21.1%) families. Overall TF families vary widely in the level of deleterious polymorphism in triads.

## Discussion

### TF retention is observed in both polyploidisation steps in wheat

In this study we found that across recurrent polyploidisation steps, wheat retains TF homoeologs more frequently than non-TF homoeologs. This is consistent with previous studies where TF retention was observed in both paleopolyploid events (>5 mya) (Li *et al*., 2016) and neopolyploid events (~7,500 to 12,500 years ago) (Zhang *et al*., 2021). Our findings agree with previous work in allotetraploid cotton *Gossypium hirsutum*, which formed through hybridisation 1-2 mya (Wendel, 1989), suggesting that TF retention is observed regardless of the time since polyploidisation. Consistent with the gene balance hypothesis, the degree of retention between different TF families was associated with both expression level and the degree of tandem duplication, demonstrating that even within a functional class, this hypothesis can make accurate predictions.

### Lack of gene expression support for homoeolog sub- or neo-functionalisation

Using the gene expression data from 15 tissues we found that overall TFs in wheat do not show sub- or neo-functionalisation at the expression or co-expression level which differs from results in other species (Liang & Schnable, 2018). For example TF duplicates formed by paleopolyploidisation events in Arabidopsis during the α, β and γ events (all > 15mya) and maize (5-12 mya) tended to have divergent expression patterns with one copy retaining ancestral expression patterns, whilst the other diverged in expression patterns (Pophaly & Tellier, 2015; Panchy *et al*., 2019). In Arabidopsis the copy with divergent expression tended to have more novel *cis*-regulatory sites, suggesting that neo-functionalisation might be happening (Panchy *et al*., 2019). One possible explanation for a lack of divergence in hexaploid wheat homoeolog expression is that the polyploidisation event is much more recent than previously studied paleopolyploidisation events. Alternatively this difference may be because wheat does not show biased genome fractionation (Juery *et al*., 2021) unlike many other studied allopolyploid species.

Although our global analysis did not show divergent patterns of expression, we found that homoeolog co-expression levels were variable between TF families. It was previously reported that a subset of triads which are dynamic in their homoeolog expression between tissues have divergent *cis*-regulation (Ramírez-González *et al*., 2018) suggesting that a small number of these changes may already be occurring in wheat. Given the highly similar expression and co-expression patterns observed in most TF families, it seems more likely that maintenance of gene dosage underlies TF retention in wheat, although sub- or neo-functionalisation of homoeolog expression may play a role in homoeolog retention in TF families that show weaker co-expression. It would require further study to establish whether *cis*-regulatory changes might explain differences in co-expression between TF families.

### Deleterious variation is reduced in TF triads indicating purifying selection

We found that hexaploid wheat TF triads have fewer deleterious missense mutations than non-TF genes. This could reflect selection against gene loss, selection against neo-functionalisation or both, i.e. purifying selection for retaining each homoeolog in its original function. Our results are consistent with Brassica allotetraploids in which TFs were enriched amongst genes without any missense mutations compared to their diploid ancestors (Zhang *et al*., 2021). However, this contrasts with paleopolyploid TF homoeologs in Brassicas which have more frequent missense mutations than other genes (Zhang *et al*., 2021). This apparent contradiction could be explained by findings from 37 angiosperm species in which TFs were enriched amongst genes which were retained in duplicate for millions of years after WGD but eventually returned to singleton status (Li *et al*., 2016). Therefore, Zhang *et al*. (2021) hypothesised that selection pressure on TFs is dynamic, with a strong purifying selection for a short period after polyploidisation (hence reduced missense mutations observed in hexaploid wheat), followed by a period with lower selection pressure once the target genes are lost through the diploidisation process. Further studies will be needed on polyploids which formed 1-5 million years ago to test this hypothesis.

### Differences between TF families

We found that TF families showed quantitative variation in their degree of diad and triad retention, degree of tandem duplication, co-expression within triads and deleterious SNP variation. While most TF families fell within a continuum of variation, the MADS-M-type family was an outlier in several analyses with the lowest percentage of genes in triads (Figure 1b) and exceptionally low co-expression levels (Figure 3) out of all 30 TF families with >10 triads. Selection to retain MADS-M-type genes appears to have been weaker than that for other TF families at both polyploidisation steps, with a gradual decrease from tetraploid to hexaploid wheat. This is consistent with previous reports that genes in the MADS-M-type family experience a high rate of birth-and-death evolution, weaker purifying selection and are less conserved between species than MADS-MIKC genes (Nam *et al*., 2004). Counter-intuitively we found that MADS-M-type triads that are retained, are highly conserved between wheat cultivars with no stop gained mutations or deleterious missense SNPs. One explanation for the contradiction of low MADS-M-type retention during polyploidisation but high conservation within hexaploid wheat cultivars could be due to their role in maintaining speciation boundaries and importance in plant reproduction (Masiero *et al*., 2011). Alternatively, the apparent high level of conservation may be due to the low number of SNPs in the MADS-M-type family included in our analysis, which is a consequence of the low level of expression of many of these genes. The MADS-M-type family contrasts strongly with the related MADS-MIKC family which behaves more similarly to other TF families and is frequently retained as triads, consistent with a previous study on the MADS-MIKC family (Schilling *et al*., 2020). While not the focus of this study, there is also likely to be extensive variation within the non-TF genes which consist of a highly heterogeneous set of genes for both function and propensity to be retained as triads.

### Implications for wheat breeding

In general we found that TF triads are retained in hexaploid wheat and have relatively few deleterious mutations, consistent with negative consequences to changing TF dosage. However, mutations in TFs which affect dosage, such as dominant mutations, have been very important in wheat breeding for their beneficial agronomic effects, for example to adapt flowering time (e.g. *VRN1* and *PPD1* (Yan *et al*., 2003; Beales *et al*., 2007)). Therefore, there is the potential to further alter gene dosage of TFs for agronomic benefit. It has been proposed that TFs with lower co-expression across tissues, termed dynamic genes (Ramírez-González *et al*., 2018), have fewer common targets (Harrington *et al*., 2020) which might release selective pressure to retain all three copies to maintain genetic balance. Therefore, one promising avenue to influence wheat phenotype by altering TF function would be to focus on TF triads with high co-expression which are more likely to have stronger phenotypic consequences if just one copy is removed. Conversely, one could focus on TF triads with low co-expression because the three homoeologs may have diverged in function, and therefore mutating one copy might lead to a phenotypic effect due to limited genetic redundancy. The recent developments in wheat functional genomics such as TILLING and gene editing now make it possible to test the effectiveness of these strategies (Krasileva *et al*., 2017; Gao, 2021).

Although the possibility to alter the sequence of one homoeolog and induce a phenotypic change in wheat is attractive, there is evidence that this will not be effective for all TFs. For example *VRN1* null mutants in a tetraploid background flower much later than wild type plants and single mutants in the A homoeolog have an intermediate flowering time, however single mutants in the B homoeolog of *VRN1* do not differ in their flowering time to WT (Chen & Dubcovsky, 2012). A similar lack of phenotype in a single mutant was observed for *NAM2* mutants which senesce at a similar time to wild type, whereas null mutants had a significant delay in senescence (Borrill *et al*., 2019). Therefore, there will still be a need for detailed functional characterisation of individual TFs, although this could be guided by predictions informed by the gene balance hypothesis.

## Acknowledgements

This work was supported by the UK Biotechnology and Biological Science Research Council (BBSRC) through grant BB/T013524/1 and the Designing Future Wheat Institute Strategic Programme (BB/P016855/1). CEBE received a BBSRC CASE Doctor Training Partnership studentship in collaboration with RAGT Seeds Ltd (BB/M01116X/1). PB also acknowledges funding from the Rank Prize New Lecturer Award. This research was also supported by the NBI Research Computing group through HPC resources and the University of Birmingham’s BlueBEAR HPC resources.

## Author contributions

PB conceived and designed the study with contributions from CEBE and RA. PB wrote the manuscript with contributions from CEBE and RA. PB carried out transcription factor annotation, triad/diad identification, tandem duplication and expression analysis. CEBE carried out SNP variant effect prediction, RA analysed nucleotide site diversity and mutation load, CEBE analysed SNP distribution in TF families. PB, CEBE and RA prepared figures and supplemental files.

## Data availability

The data that supports the findings of this study are available in the supplementary material of this article and from public repositories mentioned in the methods section. Scripts and input files are available at https://github.com/Borrill-Lab/TF_Triads.

## Supporting Information

Table S1. Genes in *Triticum aestivum* (hexaploid wheat) assigned into transcription factor families and homoeologous groups.

Table S2. *Aegilops tauschii* genes in transcription factor families.

Table S3. *Triticum urartu* genes in transcription factor families.

Table S4. Genes in *Triticum turgidum* ssp. *diccocoides* (tetraploid wheat) assigned into transcription factor families with homoeolog information.

Figure S1. Percentage of genes in triads in *Triticum aestivum* transcription factor (TF) families.

Figure S2. Percentage of genes in diads in *Triticum turgidum* ssp. *dicoccoides* transcription factor (TF) families.

Figure S3. Median expression level per TF family plotted against the percentage of the transcription factor (TF) family in triads.

Figure S4. Relationship between tandem duplication within each TF family and percentage of the transcription factor (TF) family in triads.

Figure S5. Pearson’s correlation coefficient between homoeologs across 15 tissues per transcription factor (TF) family.

Figure S6. Homoeologs in same module in 850 sample WGCNA network per transcription factor (TF) family.

Figure S7. Distribution of per-site nucleotide diversity (π) for transcription factors (TF) and background genes (non-TF).

Figure S8. Association between per-site nucleotide diversity (π) and allele frequency for transcription factors (TF) and background genes (non-TF).

## References

Avni R, Nave M, Barad O, Baruch K, Twardziok SO, Gundlach H, Hale I, Mascher M, Spannagl M, Wiebe K, et al. 2017. Wild emmer genome architecture and diversity elucidate wheat evolution and domestication. Science 357: 93.

Baker CR, Hanson-Smith V, Johnson AD. 2013. Following gene duplication, paralog interference constrains transcriptional circuit evolution. Science 342: 104–108.

Beales J, Turner A, Griffiths S, Snape JW, Laurie DA. 2007. A Pseudo-Response Regulator is misexpressed in the photoperiod insensitive Ppd-D1a mutant of wheat (Triticum aestivum L.). Theoretical and Applied Genetics 115: 721–733.

Birchler JA, Riddle NC, Auger DL, Veitia RA. 2005. Dosage balance in gene regulation: biological implications. Trends in Genetics 21: 219–226.

Birchler JA, Veitia RA. 2007. The Gene Balance Hypothesis: from classical genetics to modern genomics. The Plant Cell 19: 395–402.

Blanc G, Wolfe KH. 2004. Functional divergence of duplicated genes formed by polyploidy during *Arabidopsis* evolution. The Plant Cell 16: 1679–1691.

Borrill P, Harrington SA, Simmonds J, Uauy C. 2019. Identification of transcription factors regulating senescence in wheat through gene regulatory network modelling. Plant Physiology 180: 1740–1755.

Borrill P, Harrington SA, Uauy C. 2017. Genome-wide sequence and expression analysis of the NAC transcription factor family in polyploid wheat. G3: Genes|Genomes|Genetics 7: 3019–3029.

Borrill P, Ramirez-Gonzalez R, Uauy C. 2016. expVIP: a customizable RNA-seq data analysis and visualization platform. Plant Physiology 170: 2172–2186.

Buggs Richard JA, Chamala S, Wu W, Tate Jennifer A, Schnable Patrick S, Soltis Douglas E, Soltis Pamela S, Barbazuk WB. 2012. Rapid, repeated, and clustered loss of duplicate genes in allopolyploid plant populations of independent origin. Current Biology 22: 248–252.

Chen A, Dubcovsky J. 2012. Wheat TILLING mutants show that the vernalization gene *VRN1* down-regulates the flowering repressor *VRN2* in leaves but is not essential for flowering. PLOS Genetics 8: e1003134.

Choulet F, Alberti A, Theil S, Glover N, Barbe V, Daron J, Pingault L, Sourdille P, Couloux A, Paux E, et al. 2014. Structural and functional partitioning of bread wheat chromosome 3B. Science 345: 1249721.

Conant GC, Birchler JA, Pires JC. 2014. Dosage, duplication, and diploidization: clarifying the interplay of multiple models for duplicate gene evolution over time. Current Opinion in Plant Biology 19: 91–98.

Corey DM, Dunlap WP, Burke MJ. 1998. Averaging correlations: expected values and bias in combined Pearson rs and Fisher’s z transformations. The Journal of General Psychology 125: 245–261.

Cvijovic I, Good BH, Desai MM. 2018. The effect of strong purifying selection on genetic diversity. Genetics 209: 1235–1278.

Danecek P, Auton A, Abecasis G, Albers CA, Banks E, DePristo MA, Handsaker RE, Lunter G, Marth GT, Sherry ST, et al. 2011. The variant call format and VCFtools. Bioinformatics 27: 2156–2158.

Dubcovsky J, Dvorak J. 2007. Genome plasticity a key factor in the success of polyploid wheat under domestication. Science 316: 1862–1866.

Feldman M, Levy AA. 2012. Genome evolution due to allopolyploidization in wheat. Genetics 192: 763–774.

Force A, Lynch M, Pickett FB, Amores A, Yan YL, Postlethwait J. 1999. Preservation of duplicate genes by complementary, degenerative mutations. Genetics 151: 1531–1545.

Freeling M. 2009. Bias in plant gene content following different sorts of duplication: tandem, whole-genome, segmental, or by transposition. Annual Review of Plant Biology 60: 433–453.

Gao C. 2021. Genome engineering for crop improvement and future agriculture. Cell 184: 1621–1635.

Harrington SA, Backhaus AE, Singh A, Hassani-Pak K, Uauy C. 2020. The wheat GENIE3 network provides biologically-relevant information in polyploid wheat. G3: Genes|Genomes|Genetics 10: 3675–3686.

He F, Pasam R, Shi F, Kant S, Keeble-Gagnere G, Kay P, Forrest K, Fritz A, Hucl P, Wiebe K, et al. 2019. Exome sequencing highlights the role of wild-relative introgression in shaping the adaptive landscape of the wheat genome. Nature Genetics 51: 896–904.

Howe KL, Contreras-Moreira B, De Silva N, Maslen G, Akanni W, Allen J, Alvarez-Jarreta J, Barba M, Bolser DM, Cambell L, et al. 2020. Ensembl Genomes 2020— enabling non-vertebrate genomic research. Nucleic Acids Research 48: D689–D695.

IWGSC. 2014. A chromosome-based draft sequence of the hexaploid bread wheat (*Triticum aestivum*) genome. Science 345: 1251788.

IWGSC, Appels R, Eversole K, Stein N, Feuillet C, Keller B, Rogers J, Pozniak CJ, Choulet F, Distelfeld A, et al. 2018. Shifting the limits in wheat research and breeding using a fully annotated reference genome. Science 361: eaar7191.

Juery C, Concia L, De Oliveira R, Papon N, Ramírez-González R, Benhamed M, Uauy C, Choulet F, Paux E. 2021. New insights into homoeologous copy number variations in the hexaploid wheat genome. The Plant Genome 14: e20069.

Krasileva KV, Vasquez-Gross HA, Howell T, Bailey P, Paraiso F, Clissold L, Simmonds J, Ramirez-Gonzalez RH, Wang X, Borrill P, et al. 2017. Uncovering hidden variation in polyploid wheat. Proceedings of the National Academy of Sciences 114: E913–E921.

Langfelder P, Horvath S. 2008. WGCNA: an R package for weighted correlation network analysis. BMC Bioinformatics 9: 559.

Li Z, Defoort J, Tasdighian S, Maere S, Van de Peer Y, De Smet R. 2016. Gene duplicability of core genes is highly consistent across all angiosperms. The Plant Cell 28: 326–344.

Liang Z, Schnable JC. 2018. Functional divergence between subgenomes and gene pairs after whole genome duplications. Molecular Plant 11: 388–397.

Ling H-Q, Ma B, Shi X, Liu H, Dong L, Sun H, Cao Y, Gao Q, Zheng S, Li Y, et al. 2018. Genome sequence of the progenitor of wheat A subgenome *Triticum urartu*. Nature 557: 424–428.

Lloyd AH, Ranoux M, Vautrin S, Glover N, Fourment J, Charif D, Choulet F, Lassalle G, Marande W, Tran J, et al. 2014. Meiotic gene evolution: can you teach a new dog new tricks? Molecular Biology and Evolution 31: 1724–1727.

Luo M-C, Gu YQ, Puiu D, Wang H, Twardziok SO, Deal KR, Huo N, Zhu T, Wang L, Wang Y, et al. 2017. Genome sequence of the progenitor of the wheat D genome *Aegilops tauschii*. Nature 551: 498–502.

Lynch M, Conery JS. 2000. The evolutionary fate and consequences of duplicate genes. Science 290: 1151–1155.

Maere S, De Bodt S, Raes J, Casneuf T, Van Montagu M, Kuiper M, Van de Peer Y. 2005. Modeling gene and genome duplications in eukaryotes. Proceedings of the National Academy of Sciences 102: 5454–5459.

Masiero S, Colombo L, Grini PE, Schnittger A, Kater MM. 2011. The emerging importance of type I MADS box transcription factors for plant reproduction. The Plant Cell 23: 865–872.

McLaren W, Gil L, Hunt SE, Riat HS, Ritchie GR, Thormann A, Flicek P, Cunningham F. 2016. The Ensembl Variant Effect Predictor. Genome Biology 17: 122.

Nam J, Kim J, Lee S, An G, Ma H, Nei M. 2004. Type I MADS-box genes have experienced faster birth-and-death evolution than type II MADS-box genes in angiosperms. Proceedings of the National Academy of Sciences 101: 1910–1915.

Nowak MA, Boerlijst MC, Cooke J, Smith JM. 1997. Evolution of genetic redundancy. Nature 388: 167–171.

Ohno S. 1970. Evolution by gene duplication. Berlin Heidelberg: Springer.

Panchy NL, Azodi CB, Winship EF, O’Malley RC, Shiu S-H. 2019. Expression and regulatory asymmetry of retained *Arabidopsis thaliana* transcription factor genes derived from whole genome duplication. BMC Evolutionary Biology 19: 77.

Pophaly SD, Tellier A. 2015. Population level purifying selection and gene expression shape subgenome evolution in maize. Molecular Biology and Evolution 32: 3226–3235.

Ramírez-González RH, Borrill P, Lang D, Harrington SA, Brinton J, Venturini L, Davey M, Jacobs J, van Ex F, Pasha A, et al. 2018. The transcriptional landscape of polyploid wheat. Science 361: eaar6089.

Schilling S, Kennedy A, Pan S, Jermiin LS, Melzer R. 2020. Genome-wide analysis of MIKC-type MADS-box genes in wheat: pervasive duplications, functional conservation and putative neofunctionalization. New Phytologist 225: 511–529.

Signorell Aema. 2021. DescTools: tools for descriptive statistics. R package version 0.99.44 https://cran.r-project.org/package=DescTools.

Sim N-L, Kumar P, Hu J, Henikoff S, Schneider G, Ng PC. 2012. SIFT web server: predicting effects of amino acid substitutions on proteins. Nucleic Acids Research 40: W452–W457.

Simons KJ, Fellers JP, Trick HN, Zhang Z, Tai Y-S, Gill BS, Faris JD. 2006. Molecular characterization of the major wheat domestication gene *Q*. Genetics 172: 547–555.

Tasdighian S, Van Bel M, Li Z, Van de Peer Y, Carretero-Paulet L, Maere S. 2017. Reciprocally retained genes in the angiosperm lineage show the hallmarks of dosage balance sensitivity. The Plant Cell 29: 2766–2785.

Van de Peer Y, Fawcett JA, Proost S, Sterck L, Vandepoele K. 2009a. The flowering world: a tale of duplications. Trends in Plant Science 14: 680–688.

Van de Peer Y, Maere S, Meyer A. 2009b. The evolutionary significance of ancient genome duplications. Nature Reviews Genetics 10: 725–732.

Wendel JF. 1989. New World tetraploid cottons contain Old World cytoplasm. Proceedings of the National Academy of Sciences 86: 4132–4136.

Yan L, Loukoianov A, Tranquilli G, Helguera M, Fahima T, Dubcovsky J. 2003. Positional cloning of the wheat vernalization gene *VRN1*. Proceedings of the National Academy of Sciences 100: 6263–6268.

Zhang H, Xie J, Wang W, Wang J. 2021. Comparison of *Brassica* genomes reveals asymmetrical gene retention between functional groups of genes in recurrent polyploidizations. Plant Molecular Biology 106: 193–206.

Zheng Y, Jiao C, Sun H, Rosli HG, Pombo MA, Zhang P, Banf M, Dai X, Martin GB, Giovannoni JJ, et al. 2016. iTAK: A program for genome-wide prediction and classification of plant transcription factors, transcriptional regulators, and protein kinases. Mol Plant 9: 1667–1670.

