## Supplemental Figures for "Transcription factor retention through multiple polyploidisation steps in wheat"

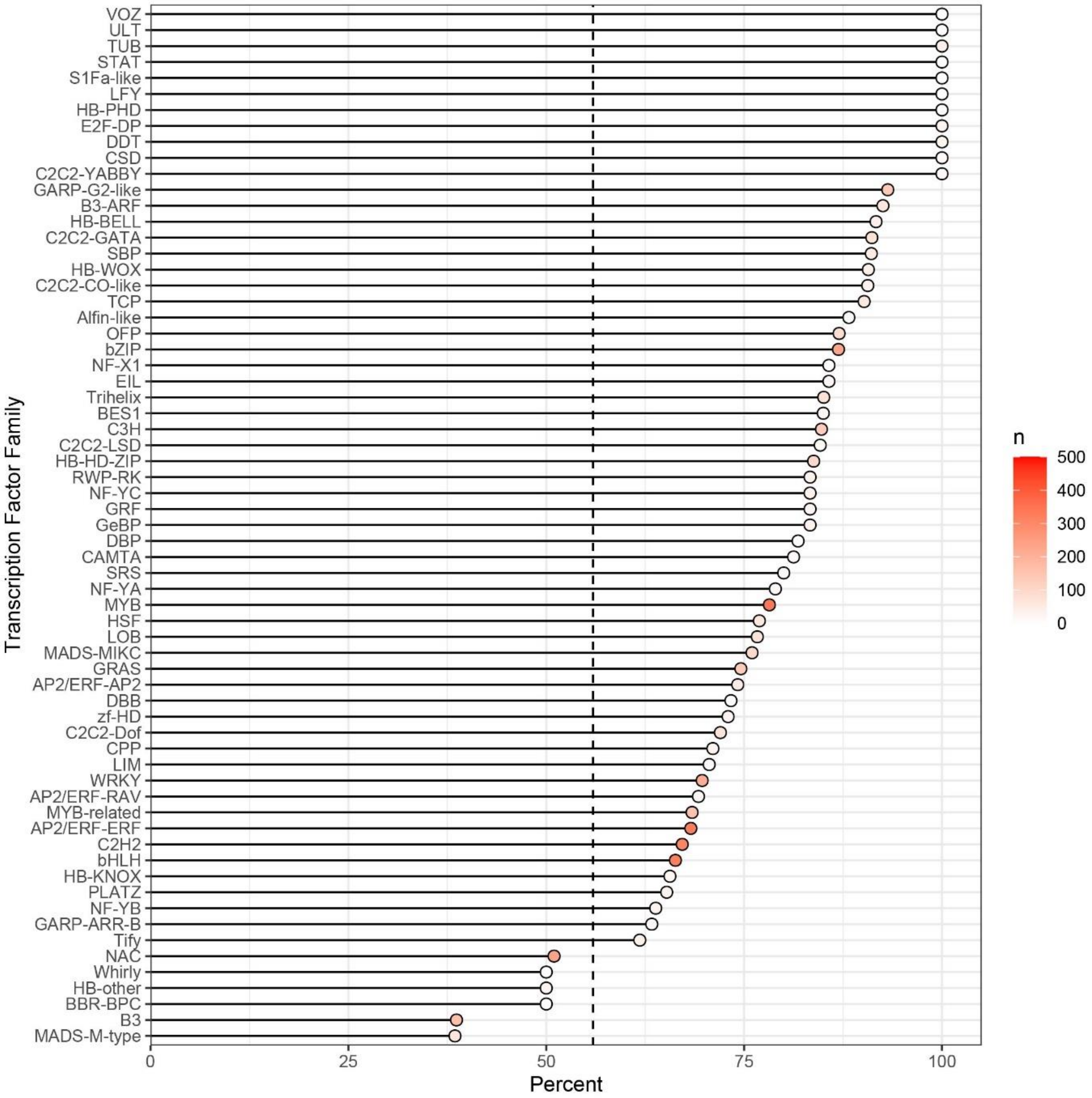


**Figure S1.** Percentage of genes in triads in *T. aestivum* transcription factor (TF) families. The dotted black line indicates the mean value for non-transcription factors. The fill colour of the dots indicates the number of genes in the TF family.

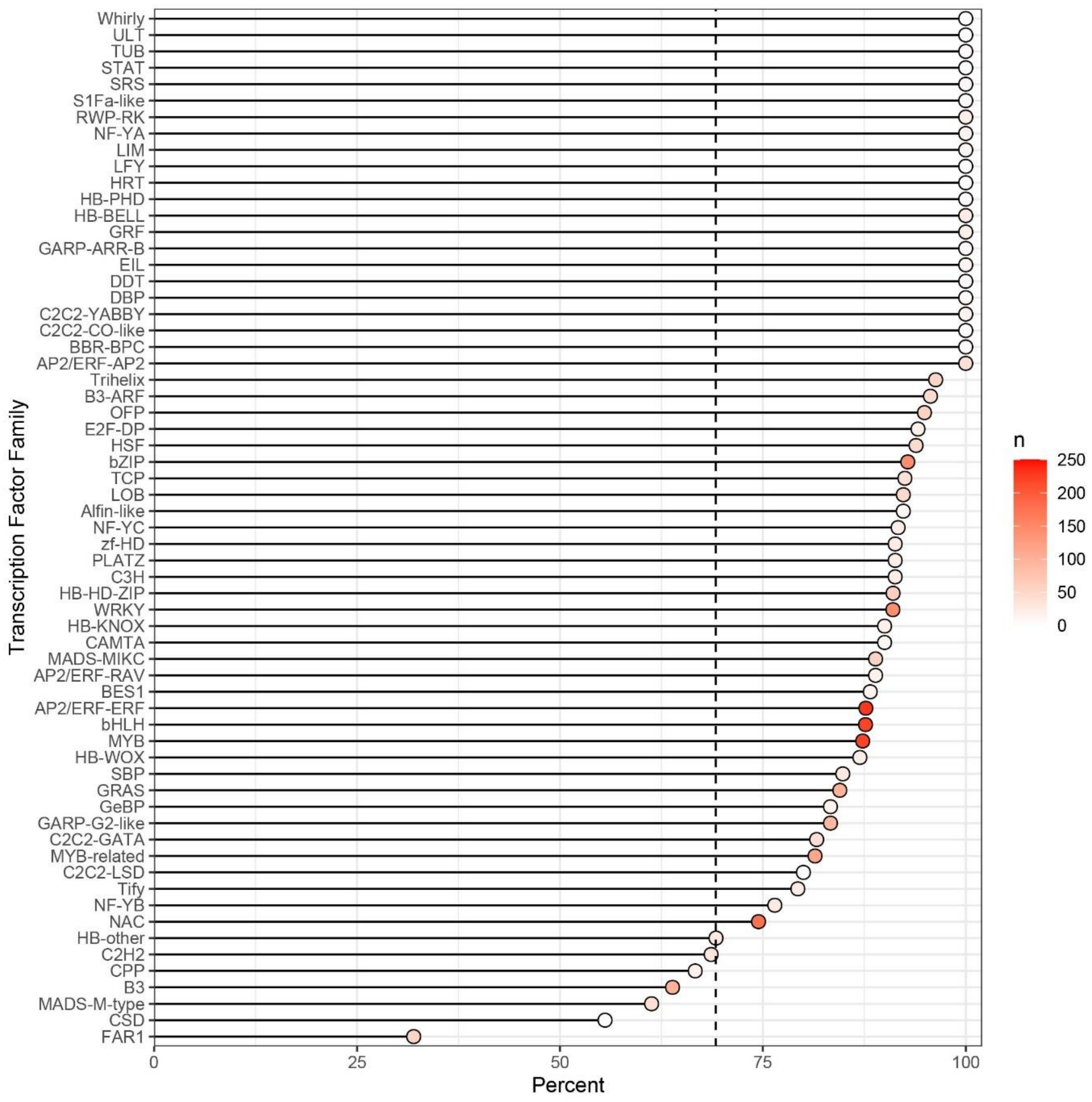


**Figure S2.** Percentage of genes in diads in *T. turgidum* ssp. *dicoccoides* T transcription factor (TF) families. The dotted black line indicates the mean value for non-transcription factors. The fill colour of the dots indicates the number of genes in the TF family.

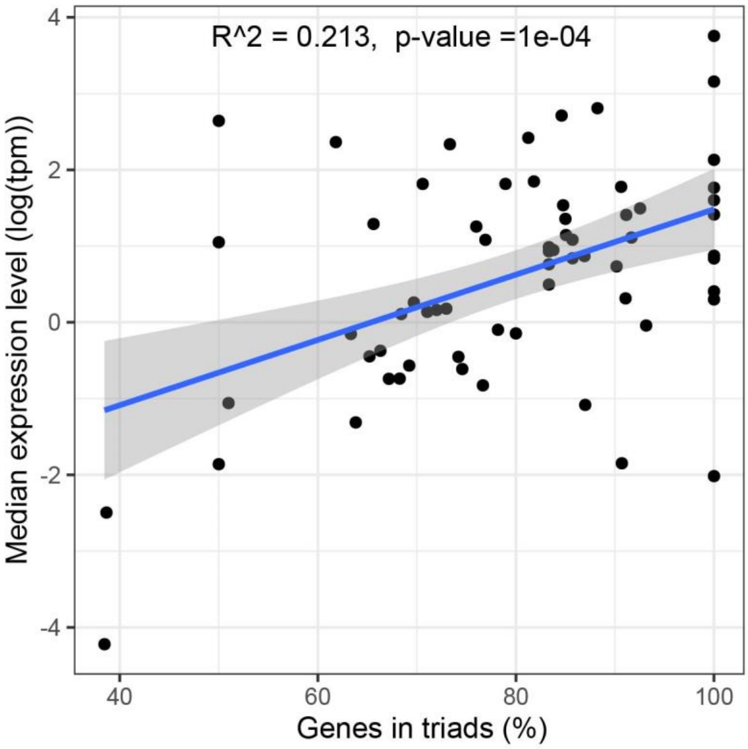


**Figure S3.** Median expression level per transcription factor (TF) family plotted against the percentage of the TF family in triads. The mean expression level of each gene was calculated using 15 tissues of Chinese Spring RNA-seq data and these gene level values were used to calculate median expression level within the TF family.

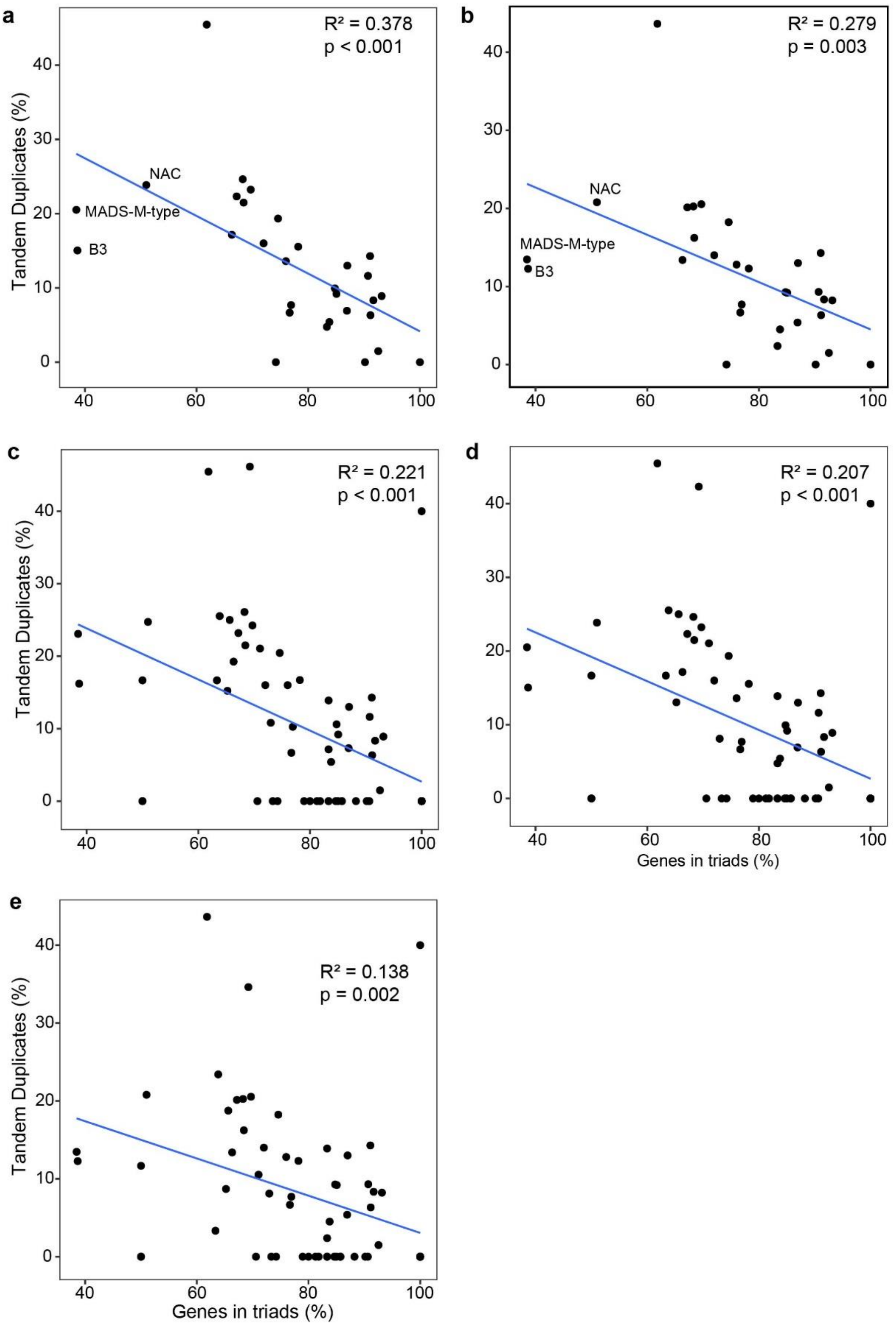


**Figure S4.** Relationship between tandem duplication within each transcription factor (TF) family and percentage of the TF family in triads. a) and b) show TF families with >30 genes in triads. c), d) and e) show all TF families. a) and d) Define tandem duplicates as up to ±2 genes away from each other (i.e. with 0 or 1 gene between duplicated TFs). b) and e) Define tandem duplicates as ±1 genes away from each other (i.e. with 0 genes between duplicated TFs). c) Defines tandem duplicates as up to ±3 genes away from each other (i.e. with 0, 1 or 2 genes between duplicated TFs).

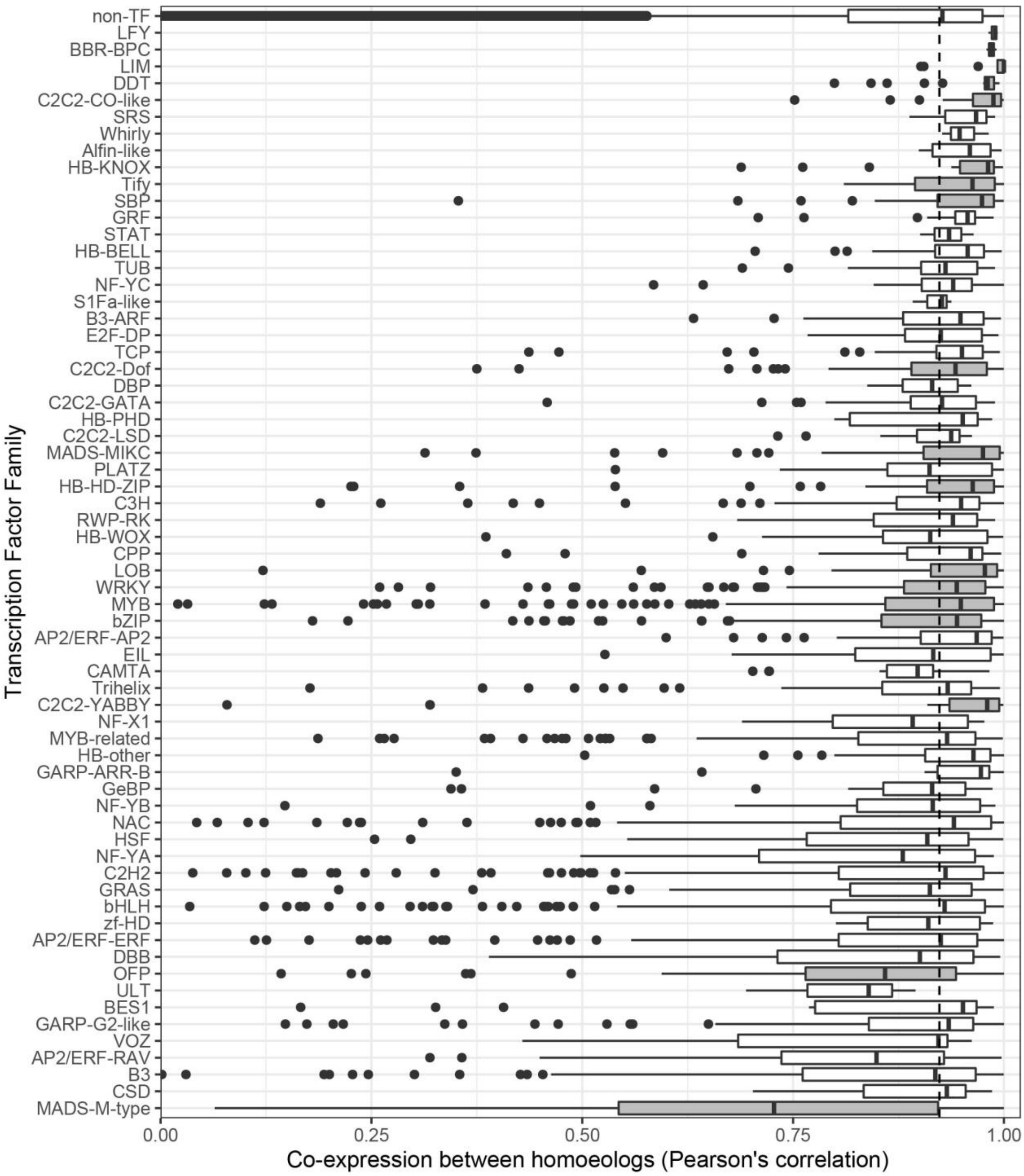


**Figure S5.** Pearson's correlation coefficient between homoeologs across 15 tissues per transcription factor (TF) family. TF families which were significantly different to non-TFs are highlighted in grey (Mann-Whitney test, $p<0.05$, FDR corrected for multiple testing). The correlation between non-TF homoeologs is shown in the top row and the dotted vertical black line represents the median value for non-TFs.

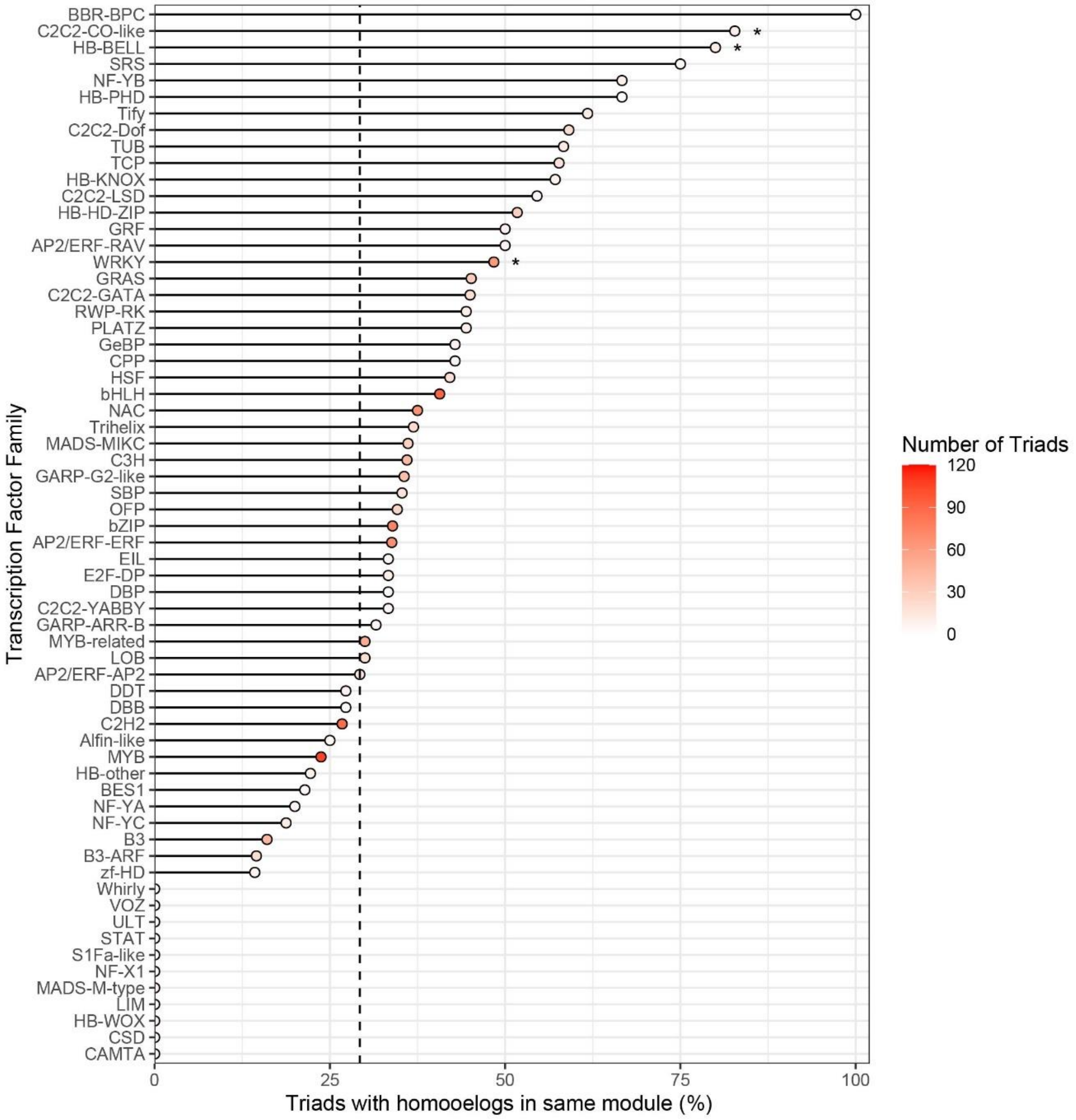


**Figure S6.** Homoeologs in same module in 850 sample WGCNA network per transcription factor (TF) family. Black dotted line represents mean value of non-TFs and asterisks (*) denote families which are statistically significant different from non-TFs

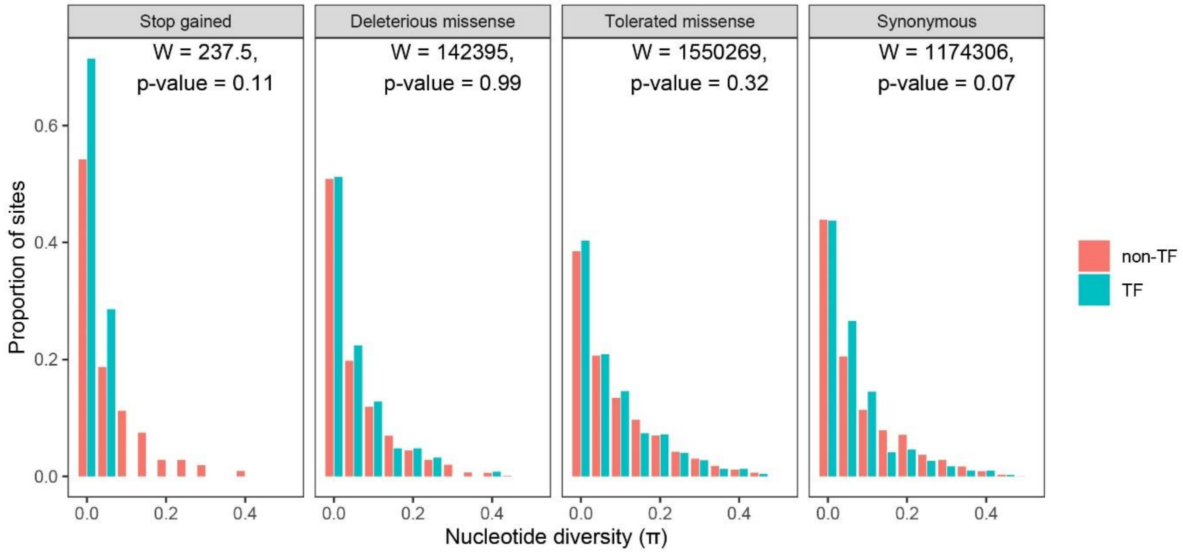


**Figure S7.** Distribution of per-site nucleotide diversity (π) for transcription factors (TF) and background genes (non-TF). A Mann-Whitney test was used to compare the TF and non-TF distributions.

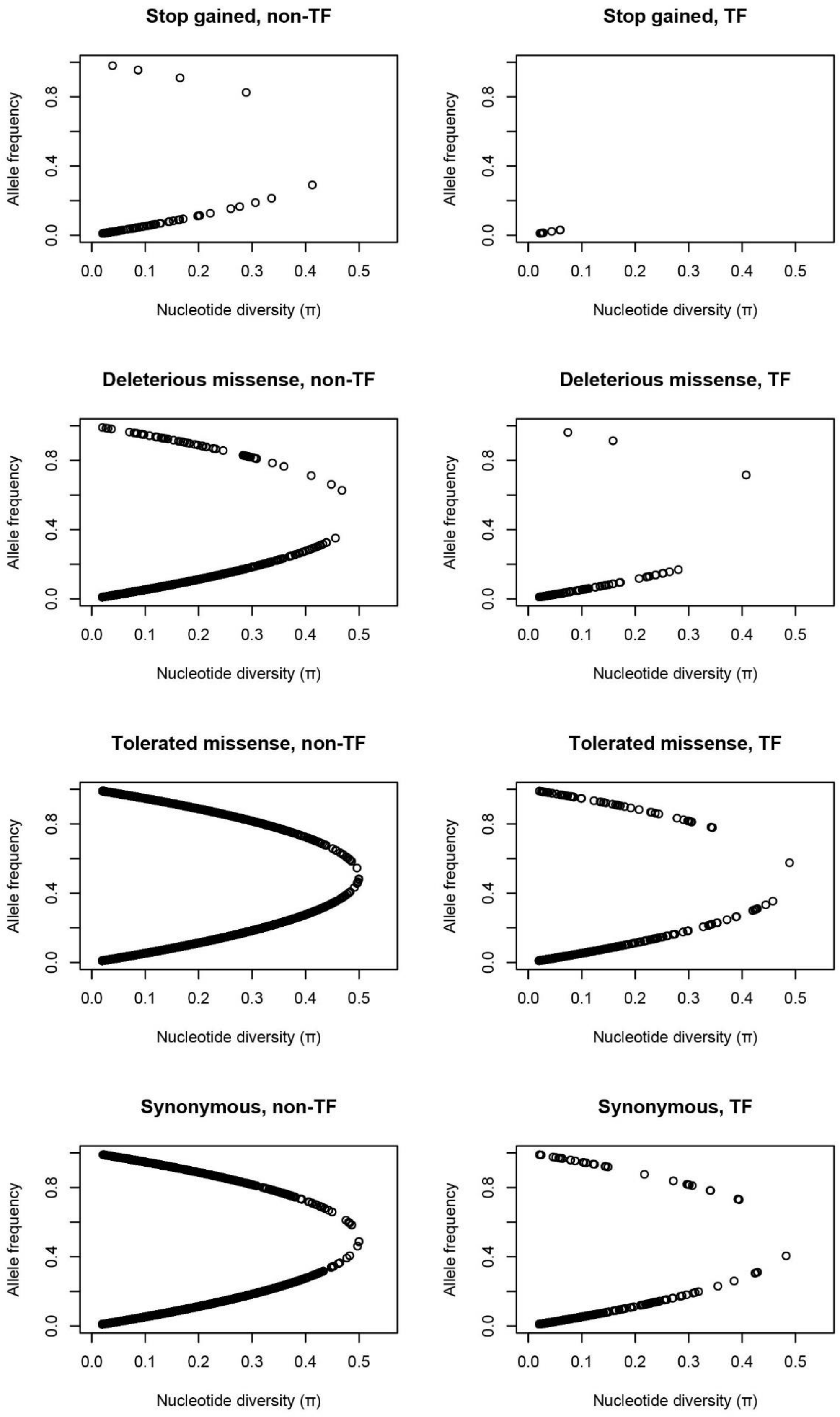


**Figure S8.** Association between per-site nucleotide diversity (π) and allele frequency for transcription factors (TF) and background genes (non-TF).
